# LONP1 is required for maturation of a subset of mitochondrial proteins and its loss elicits an integrated stress response

**DOI:** 10.1101/306316

**Authors:** Olga Zurita Rendón, Eric A. Shoubridge

## Abstract

LONP1, a AAA+ mitochondrial protease, is implicated in protein quality control, but its substrates and precise role in this process remain poorly understood. Here we have investigated the role of human LONP1 in mitochondrial gene expression and proteostasis. Depletion of LONP1 resulted in partial loss of mtDNA, complete suppression of mitochondrial translation, a marked increase in the levels of a distinct subset of mitochondrial matrix proteins (SSBP1, MTERFD3, FASTKD2 and CLPX), and the accumulation of their unprocessed forms, with intact mitochondrial targeting sequences, in an insoluble protein fraction. Depletion of LONP1 produced massive matrix protein aggregates and activated the Integrated Stress Response (ISR) pathway. These results demonstrate that LONP1 is required for maturation of a subset of its client proteins and for maintenance of mitochondrial gene expression.

## Introduction

The metazoan AAA+ proteases LONP1, CLPXP and AFG3L2 are responsible for the protein quality control of the mitochondrial matrix by degrading misfolded or oxidized polypeptides that would otherwise accumulate with deleterious consequences for mitochondrial physiology (1). Mammalian LONP1 homo-oligomerizes to form a soluble hexameric ring in which the binding and degradation of substrates takes place in an ATP-dependent manner (2–4). In addition to this proteolytic activity, it has been implicated in the maintenance of mitochondrial DNA.

In bacteria, LONP1 binds single and double-stranded DNA (5) and plays a role in the degradation of proteins involved in DNA methylation and transcription (6, 7). Deletion of Pim1, the yeast homolog of LONP1, generates major mtDNA deletions, failure to process mitochondrial-encoded mRNAs, and the accumulation of electro-dense inclusion bodies in the mitochondrial matrix (8–10). In *Drosophila melanogaster* cells, LONP1 degrades the mitochondrial transcription factor A (TFAM), a key player in the initiation of mitochondrial transcription and the major protein component of the mitochondrial nucleoid (11). In mammals, LONP1 is reported to bind only the heavy strand of the mtDNA, preferentially to G-rich sequences, and to the displacement loop (D-Loop) (12), and it plays a role in the degradation of phosphorylated TFAM (13).

The majority of nuclear-encoded mitochondrial proteins are synthesized with a N-mitochondrial targeting sequence (MTS) that is proteolytically cleaved to the mature form by the Mitochondrial Processing Peptidase (MPP), a heterodimeric enzyme composed of an alpha (MPPα) and a beta (MPPβ) subunit. The catalytic activity of mitochondrial proteases is also required for the processing of a select number of nuclear-encoded mitochondrial proteins and in the activation and/or regulation of retrograde signaling pathways. For instance AFG3L2 is responsible for the maturation of the mitochondrial ribosomal protein, MRPL32, enabling assembly of the mitochondrial ribosome (1). In the nematode, *Caenorhabditis elegans*, the accumulation of damaged proteins triggers the mitochondrial unfolded protein response (mtUPR), which is initiated in the mitochondrial matrix by the proteolytic subunit of CLPXP, CLPP, which recognizes and degrades misfolded polypeptides. This first step is followed by the activation of the bZip transcription factor ATFS-1 that accumulates in the nucleus directing transcriptional up-regulation of molecular chaperone genes (14). It is not yet clear which components of this pathway are conserved in mammals although ATF5, which can rescue an ATFS-1 deletion, has been proposed to drive a mammalian mtUPR (15). Several studies have shown that LONP1-depleted cells do not elicit this pathway (16–18), and LONP1 was shown to degrade the RNase P subunit, MRPP3A when mitochondrial proteostasis is impaired, decreasing mitochondrial translation, and thus lessening the protein folding overload (19). A recent study in which mitochondrial proteostasis was disrupted, failed to find evidence for a canonical mtUPR, but rather showed that the response to mitochondrial stress was mediated through ATF4, which directed the expression of cytoprotective genes and a reprogramming of cellular metabolism, downstream of the Integrated Stress Response (ISR) (17).

In this study we investigated the role of LONP1 in the maintenance and expression of mtDNA and in mitochondrial protein homeostasis. We show that LONP1 is essential for the maintenance of mtDNA and ribosome biogenesis. We further identify a subset of mitochondrial proteins as LONP1 substrates and show that their maturation requires the activities of both LONP1 and the MPP. Loss of LONP1 activity resulted in massive accumulation of insoluble protein aggregates in the matrix, which triggers activation of the ISR.

## Results

### Depletion of LONP1 impairs mtDNA expression

To characterize the role of LONP1 in human mtDNA metabolism we used siRNA to transiently deplete it in immortalized skin fibroblasts (LONP1-KD), and measured the effect on the levels of mtDNA, mtRNAs, and on mitochondrial protein synthesis. The levels of mtDNA were unchanged after 3 days of LONP1 depletion, but were reduced by half after 6 days (Figure 1A). There was a transient increase in the steady-state levels of all mitochondrial mRNAs tested (3 days), but after 6 days, all were substantially decreased (Figure 1B). The synthesis of all 13 mt-DNA encoded polypeptides was decreased by more than 50% after 3 days of LONPI depletion, despite the lack of change in mtDNA and the increased levels of mitochondrial mRNAs. Continued depletion of LONP1 however, completely abolished mitochondrial protein synthesis (Figure 1C), a phenotype resembling that in ϱ° (no mtDNA) cell lines.

**Figure 1.**
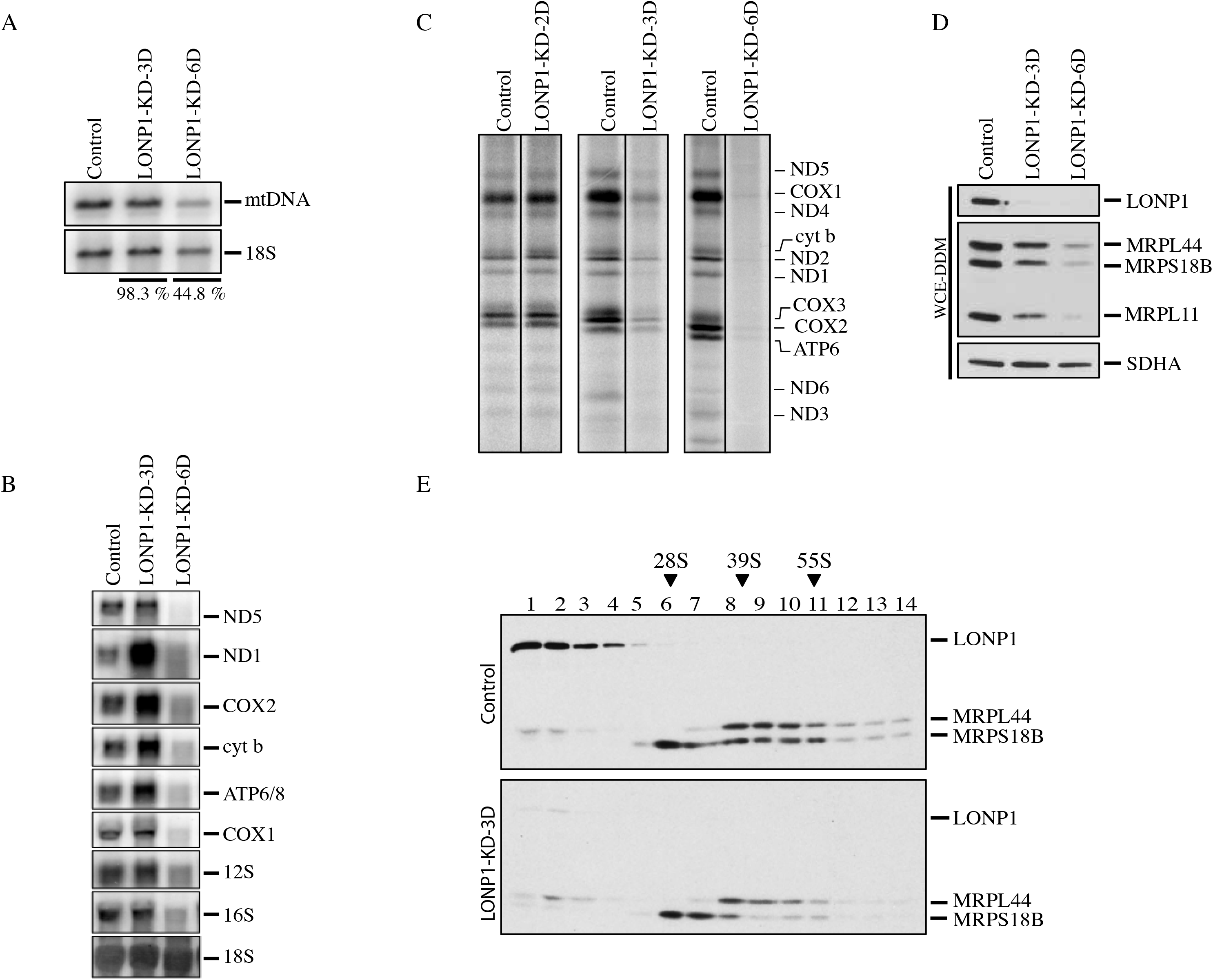
Mitochondrial gene expression in LONP1 knock-down cell lines. (A) Southern blot analysis of control and LONP1 depletion. Specific probes were used for detection of the mitochondrial DNA and the cytosolic ribosomal RNA 18S. The percentages express the levels of mtDNA in both LONP1 knock-down cell lines after normalization with the 18S (B) Northern blot analysis of control and LONP1 knock-down of 3 and 6 days. Specific probes were hybridized for detection of the mRNA subunits of complex I: ND1 and ND5, complex IV: COX1 and COX2, complex III: cyt b, complex V: ATP6/8 and the mitochondrial ribosomal RNA: 12S and 16S. The 18S cytosolic ribosomal RNA was used as loading control (C) Mitochondrial translation experiments in control and LONP1 knock-down of 2, 3 and 6 days cell lines. Mitochondrial-encoded polypeptides were pulse labeled for 60 minutes with [^35^S] methionine and cysteine. The positions of the ND subunits of complex I; COX, subunits of complex IV; cyt b, subunit of complex III; ATP, subunits of complex V are indicated on the right (D) SDS-PAGE immunodetection of LONP1 and mitochondrial ribosomal subunits (MRPs) in control and LONP1 knock-down of 3 and 6 days. Supernatants from WCE-DDM (Whole Cell Extracts with n-Dodecyl-β-D-Maltoside) were used for this analysis. SDHA is shown as a loading control (E) Sucrose density gradients were done using mitochondria from control and LONP1 knock-down of 3 days followed by immunodetection of LONP1 and MRPs. Immunoblots of MRPL44 and MRPS18B indicate the peaks for the small (28S) and large (39S) mitochondrial ribosomal subunits. 55S indicates the mitochondrial monosome.

To investigate the molecular basis for the early protein synthesis defect associated with LONP1 depletion we evaluated the assembly of the mitochondrial ribosomes. Immunoblot analysis showed a reduction in the steady-state levels of several mitochondrial ribosomal proteins (MRPs) including MRPL44, MRPS18B and MRPL11 (Figure 1D). Sucrose gradient sedimentation profiles showed a marked decrease in the fully assembled ribosome (55S), but no change in the sedimentation profile compared to control (Figure 1E). We conclude that LONP1 is required for the maintenance of mtDNA and ribosome biogenesis.

### MTERFD3 is a substrate of LONP1

To search for LONP1-interacting proteins, we immunoprecipitated endogenous LONP1 and used mass spectrometry to identify co-immunoprecipitating proteins. This analysis revealed a complex network of interactors, including the mitochondrial proteases AFG3L2, HTRA2, CLPX and YME1L1, the mitochondrial nucleoid components TFAM and SSBP1, several DNA/RNA associated proteins, the entire MICOS complex, several OXPHOS subunits and assembly factors, 7 subunits of the TIM complex and 11 mitochondrial ribosomal subunits of the large subunit (39S) including five that are part of the ribosome exit tunnel, MRPL17, MRPL22, MRPL23, MRPL39 and MRPL44 (Supplemental Figure 1).

To identify putative LONP1 substrates, we generated a fibroblast cell line (LCD) that expressed a construct coding for a catalytically dead form of LONP1 containing an S855A substitution that disrupts the catalytic dyad of LONP1 responsible for protein degradation. This substitution should not interfere with substrate binding, but would prevent subsequent proteolysis. (20). Site-directed mutagenesis was used to produce two synonymous mutations in the LONP1-siRNA target sequence, allowing us to specifically deplete endogenous LONP1 in cells expressing the inactive LONP1 (Figure 2A). Immunoblot analyses of whole cell extracts from control, LCD, LONP1-depleted cells (LONP1-KD-3D), and LCD depletion of LONP1 (LCD+LONP1-KD-3D) showed an increase in the steady-state level of LONP1 in the LCD cell line compared to control, but similar levels of expression after endogenous LONP1 was depleted (LCD+LONP1-KD-3D), confirming that we were able to express the catalytically dead LONP1, and effectively deplete endogenous LONP1 (Figure 2B). We next examined the steady state levels of MTERFD3, a protein involved in mitochondrial transcription, as a candidate substrate of LONP1 that might account for the phenotype observed in fibroblasts following LONP1 depletion. *MTERFD3* codes for a soluble protein of 43 kDa containing a mitochondrial targeting sequence (MTS) that is cleaved after translocation into the mitochondrial matrix, generating a mature 39 kDa protein (21). A unique band of 39 kDa was detected in control and LCD cell lines; however, after depletion of wild-type LONP1 mature MTERF3 was increased, and a second band of 43 kDa appeared (Figure 2B). To test whether the solubility of MTERFD3 was affected by loss of LONP1 activity, we measured the steady-state level in the soluble fraction, and found that it was reduced to nearly undetectable levels in the absence of LONP1 (Fig 2B). These results show that loss of LONP1 affects the steady state level, processing, and solubility of MTERFD3.

**Figure 2.**
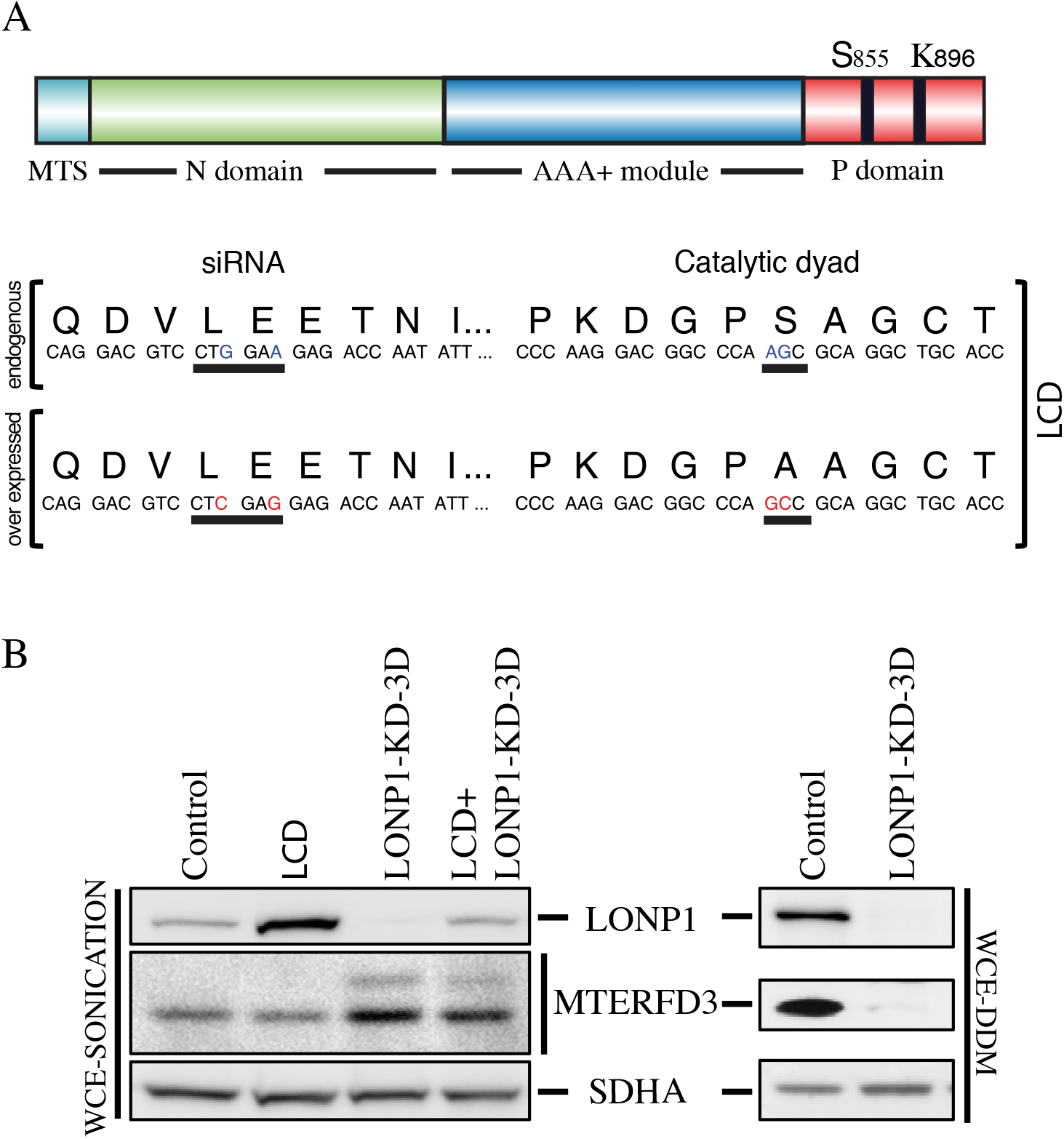
Searching for LONP1 substrates. (A) The LCD cell line overexpress a LONP1 construct containing two non-synonymous substitutions that generates a S855A mutation within the P-domain and two other synonymous mutations within the binding sequence of LONP1 siRNA that prevents it knock down (B) Western blot analysis of Whole Cell Extracts with sonication (WCE-Sonication) of indicated proteins in control, LCD, LONP1 knock-down of 3 days (LONP1-KD-3D) and LCD in which LONP1 was knocked down for 3 days (LCD+LONP1-KD-3D). SDHA is shown as loading control. Supernatants from Whole Cell Extracts with n-Dodecyl-β-D-Maltoside (WCE-DDM) were used for SDS-PAGE immunodetection of indicated proteins in control and LONP1 knock-down of 3 and 6 days. SDHA is shown as a loading control

### LONP1 is necessary for the cleavage of the mitochondrial targeting sequences for a subset of mitochondrial proteins

We next investigated the generality of the result obtained with MTERFD3 by determining the steady-state levels and solubility profile of several mitochondrial proteins in control and LCD LONP1-depleted cells (LCD+LONP1-KD-3D) after alkaline carbonate extraction (Figure 3A). We examined mitochondrial proteins covering a broad range of functions, for which we had reliable antibodies (Figure 3A).

**Figure 3.**
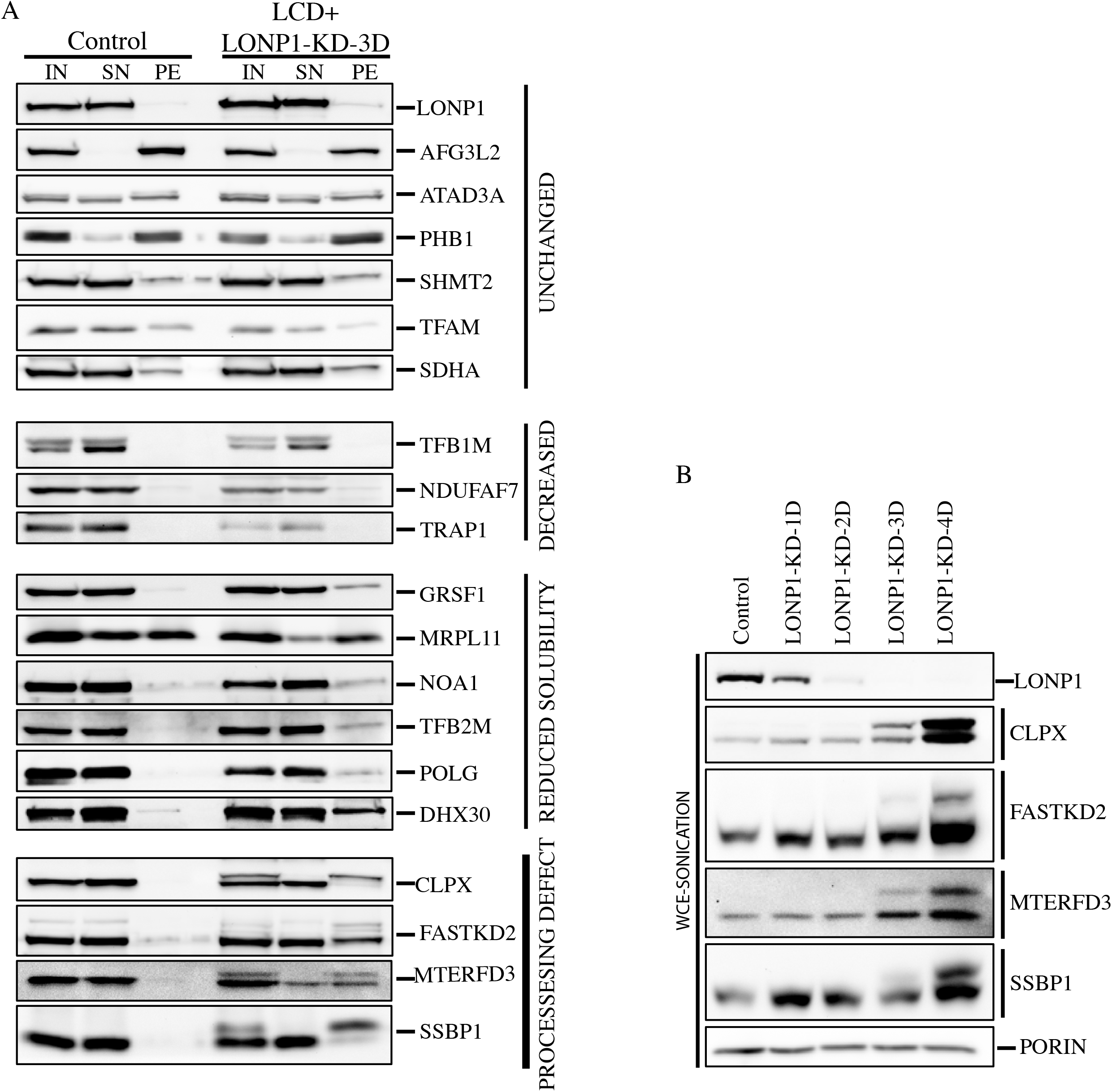
Solubilization profile of mitochondrial proteins in LONP1 knock-down cell lines. (A) Mitochondria from control and LCD in which LONP1 was knocked down for 3 days (LCD+LONP1-KD-3D) were treated with alkaline carbonate (Na_2_CO_3_). Input (IN), supernatant (SN) and pellet (PE) fractions were analyzed by SDS-PAGE (B) Whole Cell Extracts were obtained by sonication (WCE-Sonication) and used for SDS-PAGE immunodetection of indicated proteins in control and cell lines in which LONP1 was knock-down for 1,2,3 and 4 days. PORIN is shown as loading control.

The proteins could be classified into four different groups. Several, like AFG3L2 and TFAM were agnostic to the loss of LONP1 activity. A few, including the methyltransferase TFB1M, which is responsible for post-transcriptional modification of the 12S rRNA (22) were reduced, without a change in solubility. Several others, including the RNA granule proteins GRSF1 and DHX30 showed an increase in the pellet fraction. MRPL11, a structural subunit of the large ribosomal subunit (39S) was detected in the control cell line roughly equally in the supernatant and pellet fractions, but only accumulated in the pellet in LONP1-depleted cell line. Finally a forth group contained proteins, all with an apparently unprocessed mitochondrial targeting sequence, including, as expected MTERFD3, FASTKD2, an RNA granule protein that interacts with the large 16S rRNA (23, 24), the protease chaperone CLPX, and the single-stranded DNA binding protein, SSBP1.

To further characterize the processing impairment observed in SSBP1, MTERFD3, FASTKD2, and CLPX we investigated the accumulation of these proteins over 1-4 days of LONP1 depletion (Figure 3B). The loss of LONP1 correlated with increases in both the unprocessed and mature forms of these proteins, suggesting that LONP1 is necessary for both their maturation and degradation. CLPX oligomerizes with CLPP to form a mitochondrial matrix protease, and it is conceivable that its functional depletion could be the cause of the processing defect observed in LONP1-depleted cells; however, SSBP1 and MTERFD3 were correctly processed in CLPX-silenced cells (Supplemental Figure 2A).

To confirm the identity of the putatively unprocessed precursor forms of the above proteins (Figure 3A, B), we took advantage of the changes in protein solubility after LONP1 depletion. Purified mitochondria from control and LONP1-depleted cells (LONP1-KD-4D) were extracted with DDM and the pellets were solubilized with surfactant, run on SDS-PAGE gels, and silver stained. This showed a generalized increase in protein levels in the insoluble fraction of LONP1-depleted cells compared to control (Supplemental Figure 3A). Bands corresponding to FASTKD2 (~79 kDa), CLPX (~70 kDa) and MTERFD3 (~43 kDa) were excised from the gel (Supplemental Figure 3B) or analyzed in solution (SSBP1) in control and LONP-depleted cells and analyzed by mass spectrometry. As shown in Figure 4 peptides corresponding to sequences in the MTS of FASTKD2, CLPX and SSBP1 were substantially enriched in the LONPI depleted samples relative to controls. MTERFD3 is not highly expressed in human fibroblasts, and we could not detect peptides from the MTS by mass spectrometry analysis.

**Figure 4.**
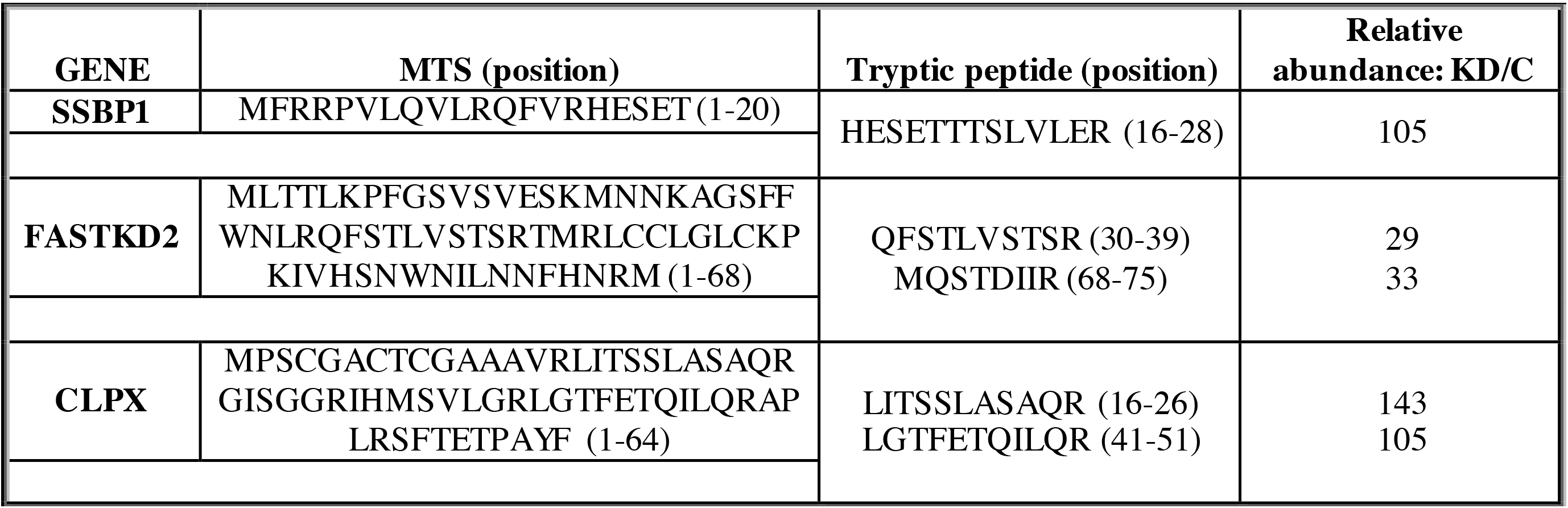
Identification of the MTS of SSBP1, FASTKD2 and CLPX. Mitochondria from control and LONP1 knock-down were treated with DDM (n-Dodecyl-β-D-Maltoside). The pellet fraction was further extracted in urea solution and ProteaseMAX™ surfactant and analyzed in solution or by SDS-PAGE and silver stained. Bands of interest were gel excised and analyzed by mass spectrometry (FASTKD2 and CLPX). The relative abundance of all peptides was calculated from the peak area of reconstructed ion chromatograms in control and LONP1 knock-down and expressed as a ratio of the knock-down over control (KD/C).

To eliminate the possibility that the processing defect observed in the above proteins was the result of deficient translocation of the precursor form to the mitochondrial matrix, we performed a Proteinase K (PK) protection assay using purified mitochondria from control and LONP1-depleted cells (Supplemental Figure 2B). This experiment showed that both the precursor and mature forms of CLPX, MTERFD3 and SSBP1 were protected from degradation at high PK concentrations, similar to the matrix protein, MRPL11.

### LONP1 and MPP are required for processing the MTS of a common subset of mitochondrial proteins

To investigate the protein composition of the insoluble fraction of LONP1-depleted cells, we suppressed LONP1 for 4 days and analyzed the pellet fraction of purified mitochondria by mass spectrometry after DDM extraction and solubilization with surfactant (Supplemental Figure 4). LONP1 depletion led to the accumulation of a very specific insoluble fraction that included the above mentioned proteins whose maturation is impaired by LONP1 depletion and proteins involved in mitochondrial translation and RNA metabolism, such as PNPT1, ELAC2, MTERFD1, ERAL1, POLRMT, NOA1, TFB2M, NOP9, GUF1, PUS1, TRUB2, TRMT10C, PTCD3, TFB1M, GFM1 amongst others. We also identified 43 mitochondrial ribosomal subunits comprising both the small (28S) and large (39S) subunits. Components of the MICOS and TIM complexes were not enriched in the pellet fraction of LONP1-depleted cells compared to controls. We further examined the MS profile of some of the proteins accumulated in LONP1-depleted cells to determine whether failure to process the MTS was a common feature. The peptides identified in 23 MRPs proteins were analyzed and we could identify peptides belonging to the predicted MTS in only MRPL43 (Supplemental Figure 3C).

PMPCB, the beta subunit of the MPP (MPPβ) was also identified in the insoluble fraction of LONP1-depleted cells suggesting that the processing defect in SSBP1, MTERFD3, FASTKD2 and CLPX could be the result of an inactive MPP. To validate the mass spectrometry analysis we used DDM to solubilize fibroblasts in control and LONP1-depleted cells (Figure 5A). Immunoblot analysis confirmed that the MPPβ subunit accumulates in the pellet fraction at 3 and 4 days of LONP1 knock-down however, the amount of MPPβ present in the soluble fraction remained comparable to control, suggesting that the depletion of LONP1 is primarily responsible for the processing defects. The MPPα subunit was only detectable in the soluble fraction in every condition, and we did not observe any changes in the processing or solubility profile of the malate dehydrogenase, MDH2, one of the validated substrates of the MPP (25).

**Figure 5.**
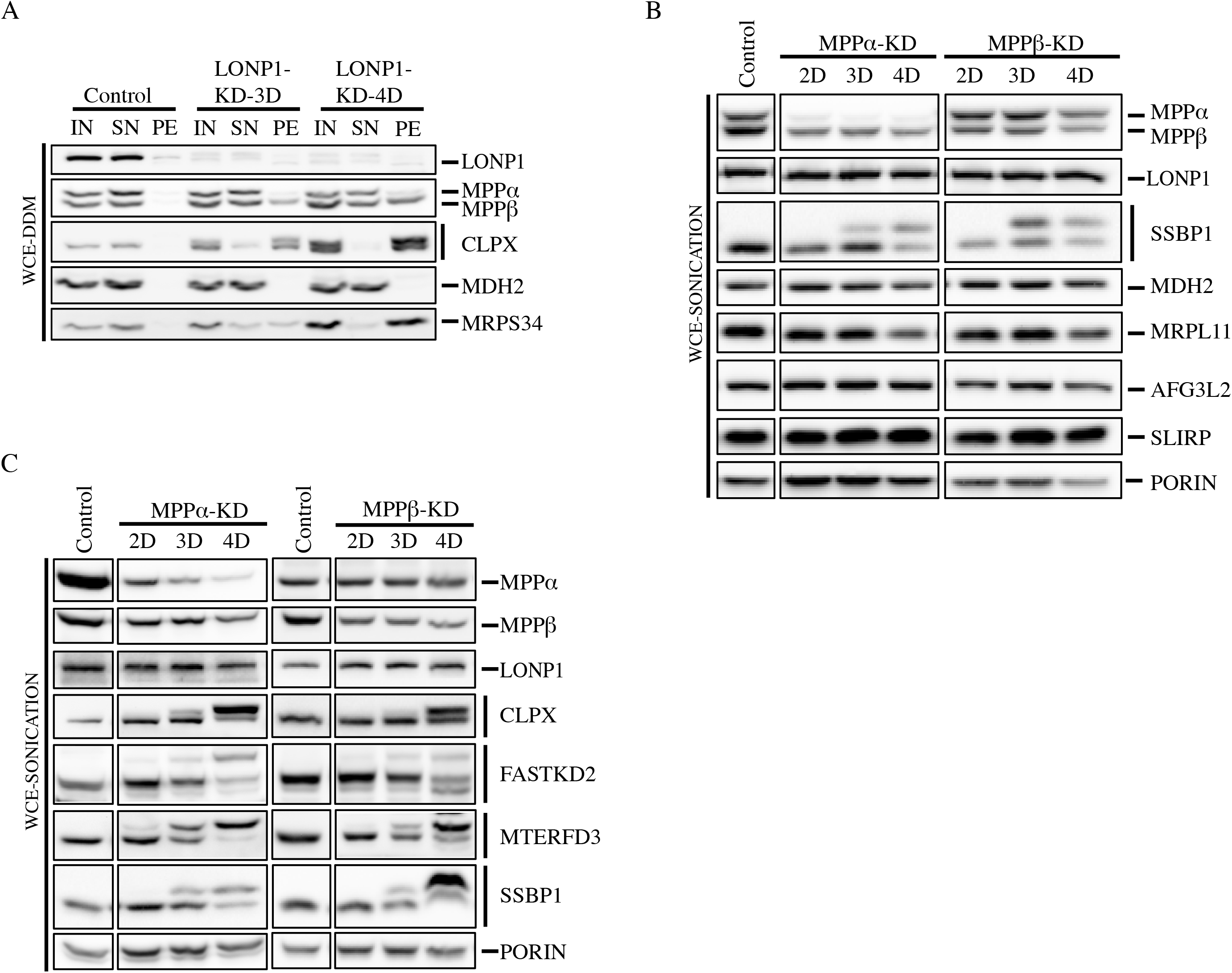
Analysis of MPP knock-down cell lines. (A) Input (IN) Supernatant (SN) and Pellet (PE) fractions from Whole Cell Extracts with n-Dodecyl-β-D-Maltoside (WCE-DDM) were used for SDS-PAGE immunodetection of indicated proteins in control and LONP1 knock-down of 3 and 4 days (B, C) Western blot analysis of Whole Cell Extracts obtained by sonication (WCE-Sonication) from control, MPPα knock down (MPPα-KD) and MPPβ knock-down (MPPβ-KD) for 2,3 and 4 days. Immunoblots show the steady state levels of indicated proteins. PORIN is shown as a loading control.

To further evaluate the role of the MPP in the processing of mitochondrial proteins we used specific siRNA constructs, MPPα-KD and MPPβ-KD, to deplete the MPPα and MPPβ subunits in a fibroblast cell line. Control and depleted cell lines were analyzed by SDS-PAGE (Figure 5B). All proteins analyzed including MDH2 were properly processed except for SSBP1. Next, we investigated if the same four proteins that showed a processing defect in LONP1-depleted cells were also affected in MPPα-KD and MPPβ-KD depletion cell lines (Figure 5C). Immunoblot analysis of CLPX, FASTKD2, MTERFD3 and SSBP1 confirmed that their processing was impaired in both MPPα and MPPβ depleted cell lines; however, in marked contrast to the results obtained in LONP1-depleted cells, MPP depletion resulted in accumulation of only the precursor, and not the mature form of the proteins.

To investigate if LONP1 and the MPP are physically associated we performed second dimension denaturing gel electrophoresis experiments (2D-SDS-PAGE) in control (Figure 6A). This experiment showed that LONP1, which is a 110 kDa protein, migrates at an apparent molecular mass of 210 kDa with a small amount smearing up to 980 kDa. The MPPα and MPPβ were present at their corresponding monomeric molecular masses of 56 and 53 kDa; however, the MPP beta subunit also accumulated at two discrete higher molecular weight complexes of about ~630 and 830 kDa, suggesting that LONP1 and MPPβ could be part of the same higher molecular weight complex. To investigate a potential interaction between LONP1 and MPPβ, we immunoprecipitated the MPP beta subunit using purified mitochondria. Immunoblot analysis showed that LONP1 was present in the elution fraction of MPPβ but not in control (Figure 6B). Mass spectrometry analysis of MPPβ immunoprecipitation confirmed the presence of LONP1 in the elution fraction (Supplemental Figure 5).

**Figure 6.**
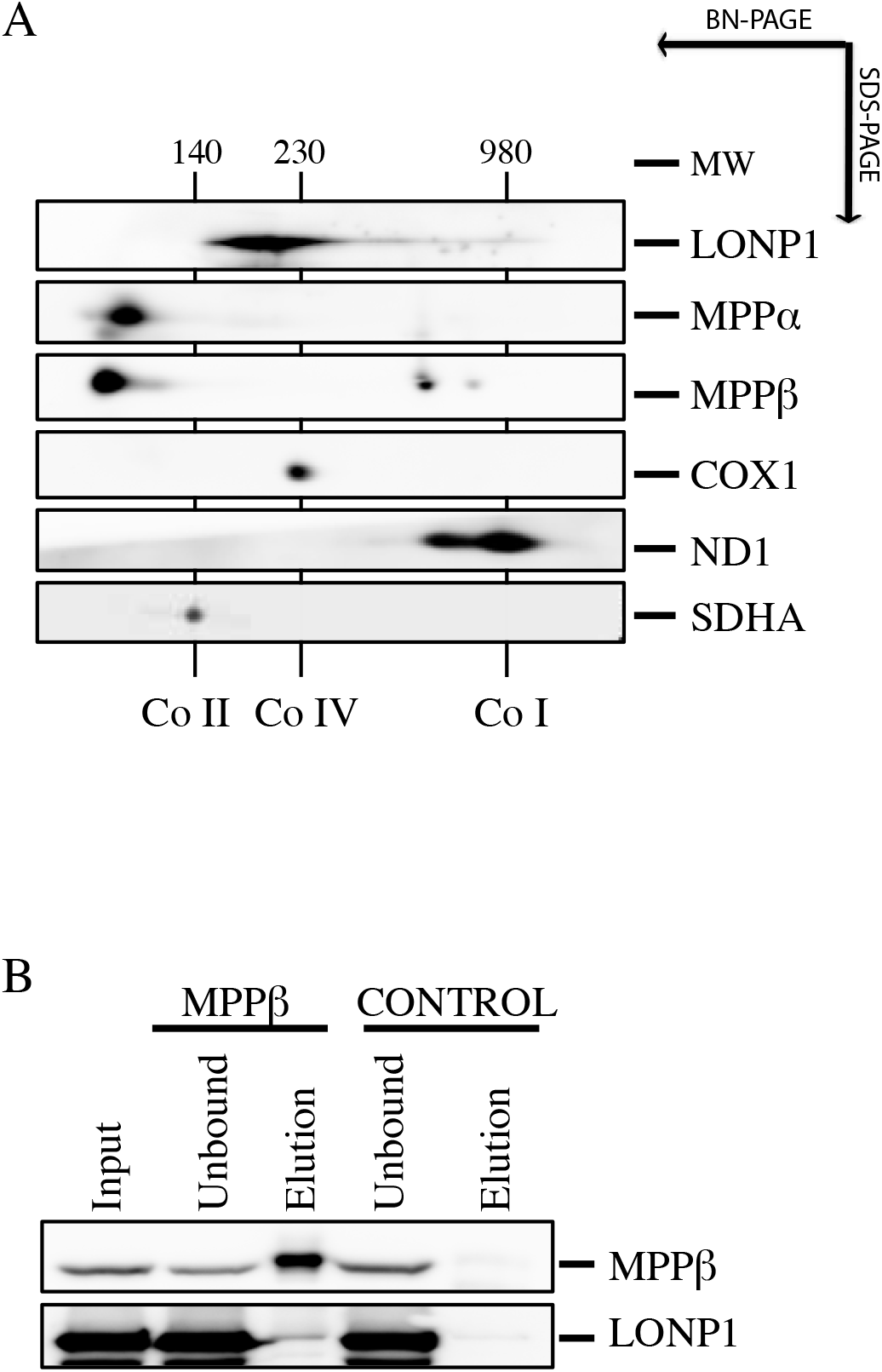
Analysis of MPP and LONP1 interaction. (A) 2D-SDS-PAGE immunodetection of LONP1 and MPP in a fibroblast control cell line. Complex I (ND1), complex IV (COX1) and complex II (SDHA) were identified with subunit specific antibodies (right). Numbers indicate the molecular weight of fully assembled OXPHOS complexes (B) MPPβ immunoprecipitation experiment was done in control mitochondria. SDS-PAGE analysis of the indicated fractions shows the steady state levels of MPPβ and LONP1.

### Depletion of LONP1 results in a massive accumulation of protein aggregates in the mitochondrial matrix

Biochemical analysis demonstrated that the solubility profile of a subset of mitochondrial proteins was altered in LONP1-depleted cells. To further study this phenomenon, we analyzed LONP1-depleted cells by electron microscopy (Figure 7). Analysis of thin sections of a control cell line showed round or ovoid mitochondria with defined cristae. After 3 days of LONP1 depletion, the cells showed a marked increase in early phagosome structures, few lysosomes and cristae-depleted mitochondria containing electro-dense aggregates. By 6 days of depletion we identified late phagosome structures, mitochondria presenting electro-dense aggregates, a more severe disruption of the cristae, and abundant cleared regions within the cytosol likely representing degraded mitochondria.

**Figure 7.**
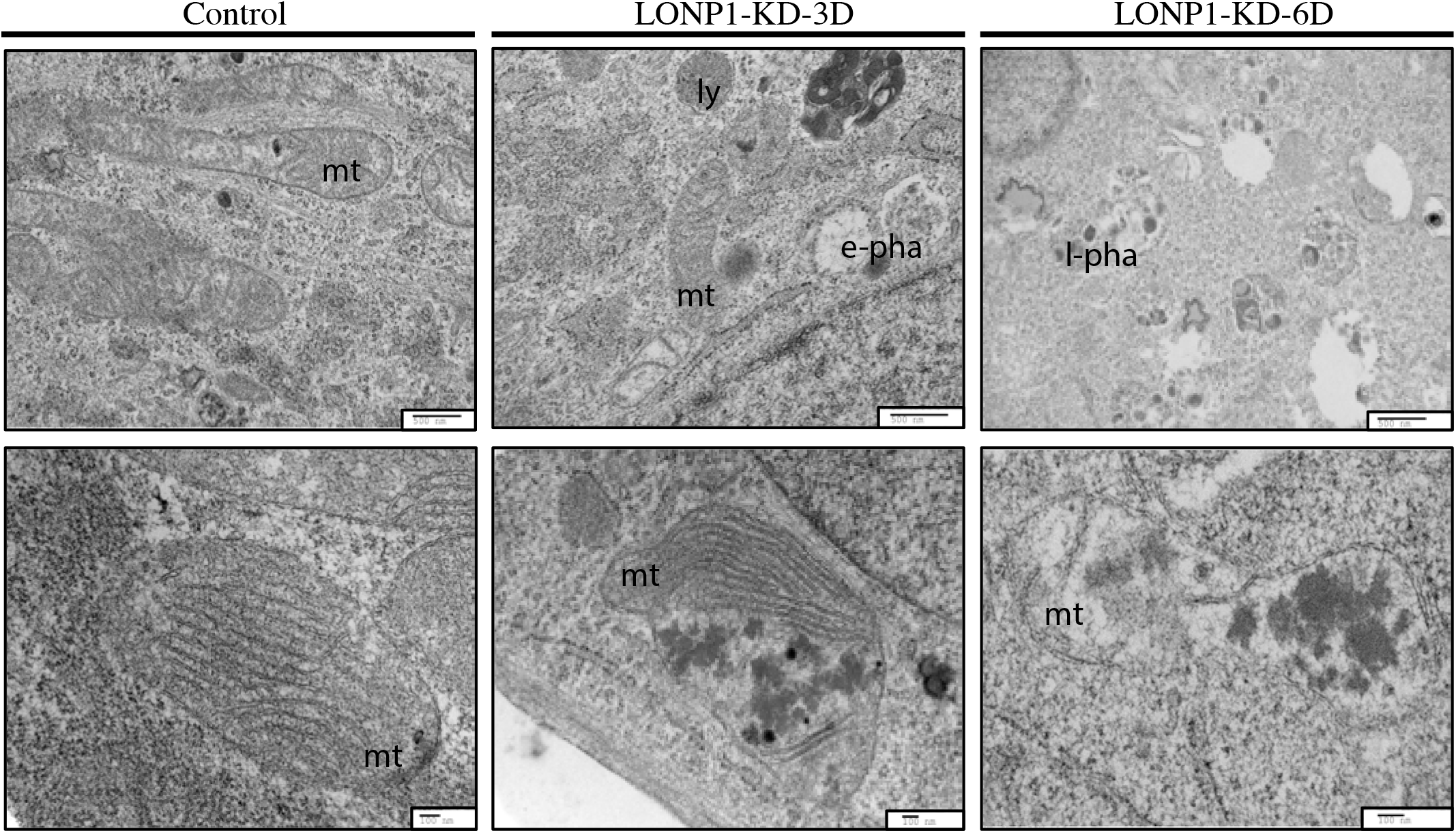
TEM analysis of LONP1 knock-down cell lines. Electron micrographs of EPON treated fibroblast sections from control and LONP1 knock-down for 3 and 6 days. mt: mitochondria, ly: lysosome, e-pha: early phagosome, l-pha: late phagosome.

### LONP1 depletion triggers an Integrated Stress Response (ISR)

When the mitochondrial protein quality control system is not sufficient to counteract the accumulation of defective polypeptides, a retrograde transcriptional pathway, the mitochondrial unfolded protein response (mtUPR), is activated to promote the synthesis of nuclear-encoded mitochondrial chaperones (15, 19, 26). To evaluate the presence of the mtUPR in the context of LONP1 depletion, we knock-down LONP1 for 2-6 days and analyzed the protein steady state levels of the CLPP protease and the chaperones HSP10, HSPA9 and HSP60 (Figure 8A). Except for CLPP, which increased slightly, all chaperones remained unaffected, even after 6 days of LONP1 depletion. We next decided to analyze the mRNA levels of the mtUPR transcriptional activator, ATF5 and several downstream components of the pathway by qRT-PCR (Figure 8B). This analysis showed that CLPP and HSPA9, significantly increased after 4 days of LONP1 depletion, however, the transcript levels of ATF5, remained unchanged, suggesting that the expression changes observed in CLPP and HSPA9 are not regulated through ATF5 activation. In contrast, mRNA quantitative analysis of the Integrated Stress Response (ISR) machinery showed that ATF4, ASNS and CHAC1 were all significantly upregulated after 4 days of LONP1 suppression. These results demonstrate that LONP1 depletion does not elicit an ATF5-driven mtUPR, but rather triggers activation of the ISR pathway (Figure 8B).

**Figure 8.**
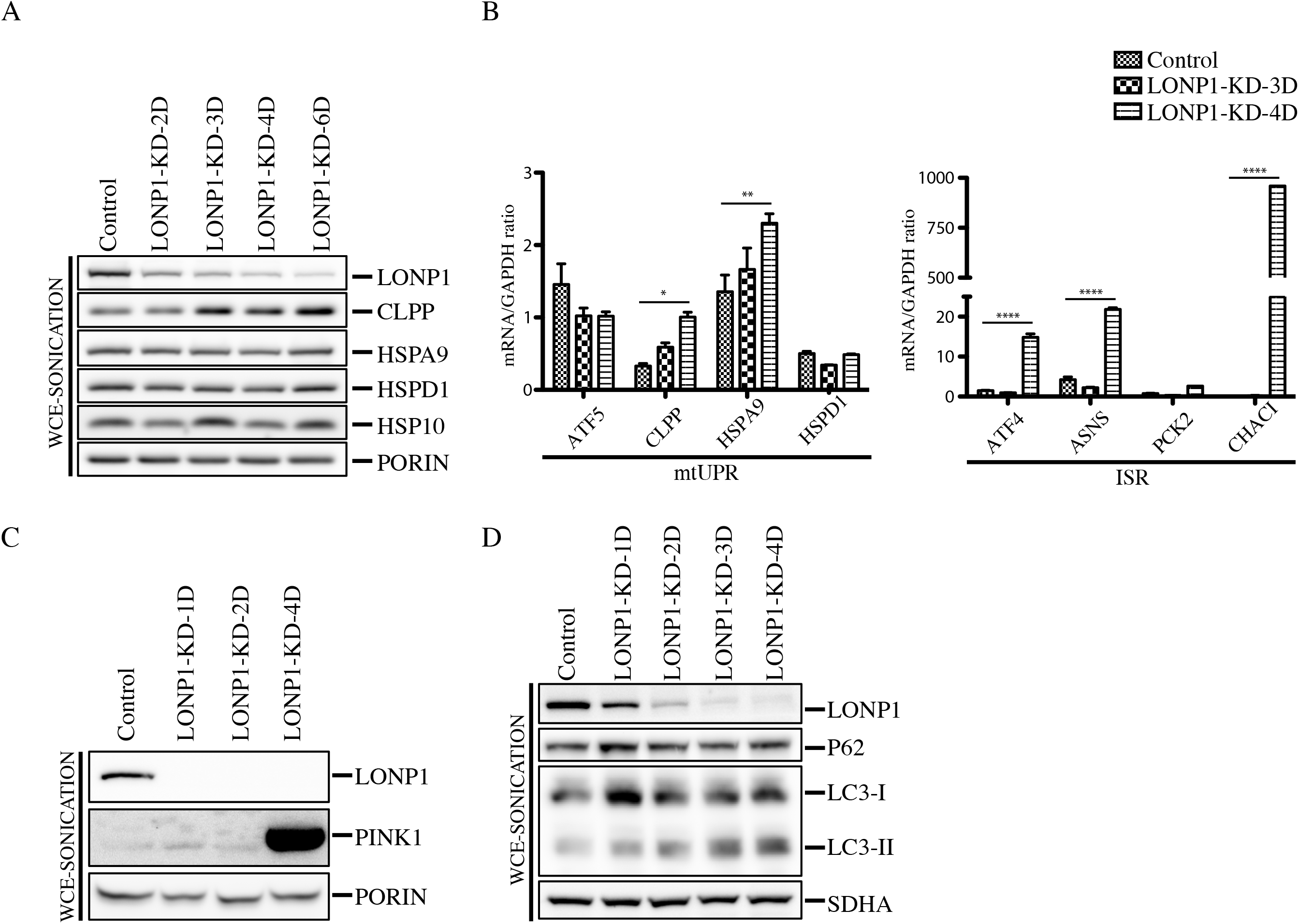
Analysis of stress responsive and mitophagy markers in LONP1 knock-down cell lines. (A) Whole Cell Extracts (WCE-Sonication) were obtained by sonication from control and LONP1 knock-down of 2-4 days and used for SDS-PAGE and immunodetection of indicated proteins. PORIN is shown as loading control (B) qRT-PCR analysis of the indicated markers of mtUPR (left) and ISR (right) was performed in control and LONP1-KD cell lines (n=3, data are mean ± s.e.m. ****P <0.0001 **P < 0.01 and *P < 0.05 (C) Whole Cell Extracts (WCE-Sonication) were obtained by sonication from control and LONP1 knock-down for 1,2 and 4 days and used for SDS-PAGE and immunodetection of indicated proteins. PORIN is shown as loading control (D) Whole Cell Extracts (WCE-Sonication) were obtained by sonication from control and LONP1 knock-down for 1-4 days and used for SDS-PAGE and immunodetection of indicated proteins. SDHA is shown as loading control.

To investigate if mitophagy was triggered in LONP1 depleted cell lines, we analyzed the steady-state level of the mitochondrial autophagy marker, PINK1, which was highly enriched in the LONP1-KD-4D cell line (Figure 8C). PINK1 is a mitochondrial Ser/Thr kinase (27) that upon mitochondrial depolarization is stabilized in the outer mitochondrial membrane where it phosphorylates ubiquitin to activate the cytosolic E3 ubiquitin ligase, PARKIN, promoting recruitment of PARKIN to mitochondria, allowing mitochondrial ubiquitination and recruitment of autophagy receptors, and subsequently mitophagy (27, 28). To further investigate the degradation pathway of damaged mitochondria after LONP1 depletion, we generated a LONP1 time course depletion followed by immunoblot analysis of whole cell extracts obtained by sonication (WCE-sonication) (Figure 8D). Immublotting of the autophagy marker LC3 revealed that the steady levels of its processed and lipidated form, LC3-II, which localizes selectively to forming and newly formed autophagosomes, increased after the first day of LONP1 knock-down. However, the autophagy receptor P62 (sequestosome 1), a ubiquitin-binding scaffold protein that directly binds LC3, and that has also been suggested to link ubiquitinated proteins to the autophagic machinery, enabling their degradation in the lysosome (29) remained unaffected.

## Discussion

Although LONP1 has long been thought of as a protease responsible for the turnover of unfolded or oxidized proteins, this study demonstrates that it also plays important roles in chaperoning/maturation mitochondrial proteins that are crucial for expression of the mitochondrial genome. We show that a decrease in mitochondrial protein synthesis, resulting from impaired ribosome biogenesis, is an early phenotype associated with loss of LONP1 function. The maturation and solubility of FASTKD2, which interacts with the 16S rRNA and plays an essential role in mitochondrial ribosome biogenesis (23, 24), is impaired when LONP1 is depleted, and this could account for the protein synthesis defect. The core nucleoid protein, SSBP1, is similarly affected, providing a plausible explanation for the defect in mtDNA maintenance.

Although LONP1 is a soluble protease of the mitochondrial matrix, immunoprecipitation experiments identified AFG3L2, YME1L1 and HTRA2 proteases, seven components of the TIM complex and the entire MICOS complex as co-immunoprecipitating proteins, all of which are intimately associated with the inner mitochondrial membrane (IMM), suggesting that LONP1 localizes near the IMM. In addition, we identified five components of the ribosome exit tunnel (MRPL17, MRPL22, MRPL23, MRPL39 and MRPL44) in this experiment. These results suggest that LONP1 functions at the boundary of IMM, interacting with a specific region of the mitochondrial ribosome. MS analysis of the pellet fraction after LONP1 suppression demonstrated that half of the structural components of the mitochondrial ribosome accumulate in insoluble protein aggregates (Supplemental Figure 4). Further evaluation of the MS data from 23 mitochondrial proteins that were found exclusively in the pellet of the LONP1-depleted cell line showed that the ribosomal subunit, MRPL43 contains peptides that are part of the predicted MTS, perhaps further contributing to the protein synthesis defect.

Depletion of the MPP (Mitochondrial Processing Peptidase) results in impaired processing of the same subset of proteins affected in LONP1 depletion, but importantly not in accumulation of the mature proteins. Mass spectrometry and immunoblot analyses of the insoluble fraction in LONP1-depleted cells identified the MPPβ subunit of the MPP; however, the alpha subunit was never detected in this fraction. A small fraction of the total amount of LONP1 and MPPβ appear to interact in a higher molecular weight complex of about 630-830 kDa. Similar to the phenotype of LONP1 depletion, the majority of the mitochondrial proteins analyzed following MPP depletion, were processed normally, including the well-characterized substrate of the MPP, MDH2. Recently, Greene et al. showed that depletion of the MPP in HEK293T cells did not affect the processing of other substrates such as UQCRFS1 (Rieske) (30, 31) or Sod2 (MnSOD)(32).

Electron microscopy analysis showed that electro-dense bodies amassed within the mitochondrial matrix of LONP1-depleted cell lines, a phenotype previously observed in yeast and humans and proposed to reflect protein aggregation (9, 33, 34). The MS analysis of the insoluble fraction of LONP1-depleted cells confirmed that the electron dense inclusion bodies of the matrix comprise a subset of mitochondrial proteins, mainly including most of the MRPs, and that the majority of these proteins were properly processed.

Determination of the components of the mammalian mtUPR have been controversial possibly due to the apparent lack of robust triggers. Despite the enormous accumulation of protein aggregates within the mitochondrial matrix in LONP1-depleted cell lines, our results show that although the expression levels of some of the components of the mtUPR, like CLPP or HSPA9, varied at the mRNA or protein level, the expression of the activator of transcription, ATF5, did not increase even after 4 days of LONP1 depletion. These results concur with several studies in which suppression of LONP1 failed to activate the canonical components of the mtUPR (17, 18). Depletion of the LONP1 homologue (Lon) in *C.elegans* similarly prevented the activation of the mtUPR (16). In contrast, instead of triggering a robust mtUPR, several components of the Integrated Stress Response, viz. ATF4, ASNS and CHAC1, which are upregulated consequent to impaired mitochondrial proteostasis to alleviate mitochondrial-related metabolic imbalances, were induced in LONP1-depleted cell lines (26).

LONP1-depleted cell lines exhibited a dramatic increase in the steady-state level of the mitophagy marker, PINK1, which is concurrent with PARKIN recruitment to fully depolarized mitochondria (27, 28). Consistently, the protein levels of the downstream autophagy marker, LC3-II, progressively accumulate with LONP1 depletion (Figure 7). The steady state levels of P62 remained unaffected, suggesting that another autophagy receptor is responsible for the binding to the ubiquitinated cargo. Interestingly, the LC3-II protein levels increased after 24 hours of LONP1 depletion; however, the PINK1-PARKIN pathway was only active in LONP1-4D-KD cell lines. It is possible that the initial pathway of mitochondrial degradation in the context of LONP1 depletion is P62/PARKIN-independent as recently proposed by Lazarou M. et al. (35).

Together our results demonstrate that the human protease LONP1 is required for the maturation and degradation of a subset of mitochondrial matrix proteins with key roles in the expression of the mitochondrial genome. Failure to process these substrates presumably prevents their proper folding, resulting in their accumulation in insoluble protein aggregates, ultimately triggering an ISR and mitophagy. Why a specific subset of mitochondrial matrix proteins require LONP1 activity for proper processing remains an outstanding question. It is possible that the tertiary structure of immature SSBP1, MTERFD3, FASTKD2 and CLPX differ from the “canonical” features required for processing by MPP alone, but this will require further investigation.

## Materials and Methods

### Cell culture

Primary cell lines were established from subject skin fibroblasts and immortalized by transduction with a retroviral vector expressing the HPV-16 E7 gene plus a retroviral vector expressing the catalytic component of human telomerase (htert) (36). Fibroblasts and 143B cell lines were grown at 37 °C in an atmosphere of 5% CO_2_ in high-glucose Dulbecco’s modified Eagle medium (DMEM) supplemented with 10% fetal bovine serum.

### Mitochondrial isolation

Fibroblasts were resuspended in ice-cold SET buffer: 250 mM sucrose/10 mM Tris-HCl/1 mM EDTA (pH 7.4) supplemented with complete protease inhibitors (Roche) and homogenized with 10 passes through a pre-chilled, zero clearance homogenizer (Kimble/Kontes). The homogenized cellular extract was then centrifuged twice for 10 minutes at 600g to obtain a postnuclear supernatant. Mitochondria were pelleted by centrifugation for 10 minutes at 10,000g, and washed once in the same buffer.

### LONP1 overexpression

cDNA from LONP1 was amplified by OneStep RT-PCR (QIAGEN) and cloned into pBABE-Puro with the Gateway Cloning Technology (Invitrogen). The accuracy of the clones was tested by automated DNA sequencing. Direct mutagenesis of LONP1 was done using the site direct mutagenesis lightning kit (Stratagene). Retroviral constructs were transfected into the Phoenix packaging cell line using the HBS/Ca_3_(PO_4_)_2_ method. Control fibroblasts were infected 48 hours later by exposure to virus-containing medium in the presence of 4 mg/ml of polybrene as described (http://www.stanford.edu/group/nolan/protocols/pro_helper_dep.html).

### RNAi knock-down

For the knockdown experiments of LONP1, CLPX and MPP, a Stealth RNA interference (RNAi) duplex construct was used as follows:

LONP1: GGACGUCCUGGAAGAGACCAAUAUU

CLPX: AAUAUCUUCGCCUACAUAUCCAGCC

MPPαa: GCAUGUUCUCCAGGCUCUACCUCAA

MPPαb: GAGCCAAGACGCAGCUGACAUCAAU

MPPβa: GCAGCUUGAUGGUUCAACUCCAAUU

MPPβb: CAGCUCACUUGUCAUGGCAAUCUUU

siRNA constructs were designed through the BLOCK-iT™ RNAi Express website: http://rnaidesigner.invitrogen.com/rnaiexpress/rnaiExpress.jsp?CID=FL-RNAIEXPRESS

Indicated cell lines were transfected following the Lipofectamine™ RNAiMAX protocol. Briefly, lipofectamine and RNA constructs were diluted in Opti-MEM Reduced Serum Medium. BLOCK-iTAlexa Fluor Red Fluorescent Oligo was used to assess the transfection efficiency (transfection performed in parallel with the other stealth constructs) and as a mock oligo transfection control (All reagents from Invitrogen).

### Southern blot

DNA was isolated from LONP1 siRNA knocked-down and control fibroblasts using the genElute mammalian genomic DNA miniprep kit (Sigma). Five micrograms of total DNA were PvuII digested overnight at 37 °C. Samples were run on a 1% agarose/TBE gel follow by 20 minutes incubations at room temperature for depurination, denaturation and neutralization. Next, gel was transfer to a nylon membrane. mtDNA and 18S probes were labeled with [^32^P]-dCTP (GE Healthcare) using the MegaPrime DNA labeling kit (GE Healthcare). Hybridization was conducted according to the manufacturer’s manual using ExpressHyb Hybridization Solution (Clontech) and the radioactive signal was detected using the Phosphoimager system. Quantitative analysis was done using the ImageQuant 5.2 software.

### Northern blot

RNA was isolated from LONP1 siRNA knocked-down and control fibroblasts using the RNeasy Kit (QIAgen). Ten micrograms of total RNA were separated on a denaturing MOPS/formaldehyde agarose gel and transferred to a nylon membrane. ND1, ND5, COX1, COX2, ATP6/8, cyt b, 12S, 16S and 18S probes were labeled with [^32^P]-dCTP (GE Healthcare) using the MegaPrime DNA labeling kit (GE Healthcare). Hybridization was conducted according to the manufacturer’s manual using ExpressHyb Hybridization Solution (Clontech) and the radioactive signal was detected using the Phosphoimager system.

### Pulse labeling of mitochondrial translation products

In vitro labeling of mitochondrial translation was performed as previously described (37). Briefly, cells were labeled for 60 minutes at 37°C in methionine and cysteine-free DMEM containing 200 *μ*Ci/ml [35S]methionine and 100 *μ*g/ml emetine and chased for 10 minutes in regular DMEM. Protein extraction was done in labeled cells by resuspension in loading buffer containing 93 mM Tris-HCl, pH 6.7, 7.5% glycerol, 1% SDS, 0.25 mg bromophenol blue/ml and 3% mercaptoethanol. Samples were sonicated for 5 seconds and run on 15–20% polyacrylamide gradient gels. The labeled mitochondrial translation products were detected by direct autoradiography on a Phosphoimager.

### SDS-PAGE analysis

Whole Cell Extracts with n-Dodecyl-β-D-Maltoside (WCE-DDM) were prepared by solubilization in approximately 1:5 ratio with extraction buffer (1.5% DDM) for 30 minutes on ice followed by centrifugation at 20 000*g* for 20 minutes at 4°C. The soluble fraction was mixed at a 1:1 ratio with 2X Laemmli Sample Buffer. The same protocol of extraction was used for the separation of the soluble (SN) and insoluble (PE) fraction of LONP1-KD and MPP-KD cell lines. Whole cell extracts by sonication (WCE-sonication) was done by resuspension of pelleted cells in loading buffer containing 93 mM Tris-HCl, pH 6.7, 7.5% glycerol, 1% SDS, 0.25 mg bromophenol blue/ml and 3% mercaptoethanol. Next, samples were sonicated for 5 seconds. WCE-DDM and WCE-sonication samples were loaded onto a 12% acrylamide:bisacrylamide (29:1) gel. Subsequent immunoblotting was done with indicated antibodies.

### Antibodies

The following antibodies were obtained as follows: anti-LONP1 (kindly provided by Carolyn Suzuki), anti-ACTIN (MitoSciences), anti-SDHA (Abcam), anti-PORIN (EMD Calbiochem), anti-PHB1 (Abcam), anti-LRPPRC (in house), anti-SLIRP (Abcam), anti-MRPL11 (Sigma-Aldrich), anti-MRPS18B (Proteintech Group), anti-MRPL44 (Sigma-Aldrich), anti-MTERFD3 (Sigma-Aldrich), anti-TFAM (a kind gift of Alexandra Trifunovic), anti-TFB1M (Sigma-Aldrich), anti-TFB2M (Abgent), anti-AFG3L2 (kindly provided by T. Langer), anti-GRSF1 (Sigma-Aldrich), anti-ATAD3A (Abcam), anti-SHMT2 (Abcam), anti-NDUFAF7 (in house), anti-NOA1 (Sigma), anti-POLG (Abcam), anti-CLPX (Abcam), anti-SSBP1 (Novus biological), anti-DHX30 (Abcam), anti-FASTKD2 (Santa cruz), anti-MFN2 (Cell signaling), anti-SCO1 (in house), anti-MPPα (Sigma), anti-MPPβ (proteintech), anti-MDH2 (proteintech), anti-COX1 (abcam), anti-ND1 (kindly provided by A. Lombes), anti-LC3 (Novus biologicals), anti-P62 (BD biosciences), anti-YME1L1 (Proteintech), anti-CLPP (Proteintech), anti-HSP10 (Santa cruz), anti-HSPA9 (Proteintech), anti-PINK1 (Novus biological)

### Sucrose density gradient

Mitochondria were extracted in lysis buffer containing 260 mM sucrose, 100 mM KCl, 20 mM MgCl2, 10 mM Tris-Cl, pH 7.5, 1% Triton X-100 and 5 mM β-mercaptoethanol, supplemented with complete protease inhibitors without EDTA, on ice for 20 minutes. Lysates were centrifuged for 45 minutes at 9400g at 4°C before loading them on 1 ml of 10-30% discontinuous sucrose gradient (50 mM Tris-Cl, 100 mM KCl, 10 mM MgCl2). Loaded gradients were centrifuged at 32,000 rpm for 130 minutes in a Beckman SW60-Ti rotor. After centrifugation, 14 fractions were collected and analyzed by SDS-PAGE.

### Immunoprecipitation

Mitochondria from control (LONP1 and MPPβ immunoprecipitations) cell lines were DSG (1 mM Disuccinimidyl glutarate) cross-linked for 90 minutes on ice prior to protein extraction. MPPβ immunoprecipitation experiment was not cross-linked. One miligram of mitochondria was extracted on ice in 250 μl of lysis buffer (50 mM HEPES buffer, pH 7.6, 150 mM NaCl, 1% taurodeoxycholate), supplemented with complete protease inhibitors (Roche), for 45 minutes. The protein extract was centrifuged at 25,000g at 4°C, for 40 minutes. Supernatant was then incubated overnight with naked magnetic Dynabeads for preclearance; in parallel Dynabeads were coated to the appropriate antibody (LONP1 or MPPβ) according to the manufacturer’s instructions. The incubation of the protein extract with antibody coated beads was carried out overnight at 4°C. Bound protein was elute with 400 μl of 0.1M glycine pH 2.5 with 0.5% DDM at 45°C followed by TCA precipitation. Elutions were trypsin digested and analyzed by mass spectrometry using a LTQ Orbitrap Velos (ThermoFisher Scientific, Bremen, Germany).

### Sodium carbonate solubilization

Purified mitochondria from control and LCD+LONP1-KD-3D fibroblasts were extracted with 100 mM Na_2_CO_3_ pH 11.5 on ice for 30 minutes and centrifuged at 65,000 rpm to separate soluble (SN) and membrane (PE) fractions. Pellet (PE) fraction was washed twice with water prior to SDS-PAGE analysis. Subsequent immunoblotting was done with indicated antibodies.

### Surfactant treatment and silver stained analysis

Mitochondria from control, LONP1-KD-4D and LONP1-KD-6D were resuspended in approximately 1:5 ratio with extraction buffer (1.5% DDM) for 30 minutes on ice followed by protein quantification by Bradford assay. Equal amounts of protein from control and knock-down cell lines were centrifuged at 20 000g for 20 minutes at 4°C. The supernatant fraction was collected and the pellet was rinse in acetone followed by resuspension in 8M urea solution and 0. 2% ProteaseMAX™ surfactant. Samples were incubated at room temperature for 60 minutes followed by centrifugation at 16 000g for 10 minutes. Next, supernatants were directly send for mass spectrometry analysis and/or run by SDS-PAGE followed by silver staining; briefly gels were fixed in 50% methanol, 10% acetic acid solution for 30 minutes, reduced in 0.02% sodium thiosulfate for 2 min and stained with 0.2 % silver nitrate for 30 minutes. Developing was done with a solution containing 0.05% formalin, 3% sodium carbonate and 0.0004% sodium thiosulfate. Bands of interest were gel excised, digested with trypsin or glu-c and analyzed by mass spectrometry using a LTQ Orbitrap Velos (ThermoFisher Scientific, Bremen, Germany).

### Mass spectrometry analysis

The in-gel digestion protocol is based on the results obtained by (38). Bands of interest were gel excised and all volumes were adjusted according to the volume of gel pieces. Gel pieces were washed with water for 5 minutes, destained twice with the destaining buffer (100 mM sodium thiosulfate, 30 mM potassium ferricyanide) for 15 minutes. This step was followed by 5 minutes wash with 50 mM ammonium bicarbonate and then dehydrated with acetonitrile. Next, reduction buffer (10 mM DTT, 100 mM ammonium bicarbonate) was added and incubated for 30 minutes at 40°C, followed by a 20 minutes incubation at 40°C in alkylation buffer (55 mM iodoacetamide, 100 mM ammonium bicarbonate). Gel pieces were dehydrated and washed at 40°C by adding acetonitrile for 5 minutes and then dried for 5 minutes at 40°C and then rehydrated at 4°C for 40 min with the trypsin solution (6 ng/*μ*l of trypsin or glu-c sequencing grade from Promega, 25 mM ammonium bicarbonate). The concentration of trypsin or glu-c was kept low to reduce signal suppression effects and background originating from autolysis products when performing LC-MS/MS analysis. Protein digestion was performed at 58°C for 1 hour and stopped with 1% formic acid/2% acetonitrile. Supernatant was transferred into a 96-well plate and peptides extraction was performed with two 30 minutes extraction steps at room temperature using the extraction buffer (1% formic acid/50% acetonitrile). All peptide extracts were pooled into the 96-well plate and then completely dried in vacuum centrifuge. The plate was sealed and stored at 20°C until LC-MS/MS analysis. Prior to LC-MS/MS, peptide extracts were re-solubilized under agitation for 15 minutes in 11 *μ*l of 0.2% formic acid and then centrifuged at 2000 rpm for 1 minutes.

The LC column was a C18 reversed phase column packed with a high-pressure packing cell. A 75 *μ*m i.d. Self-Pack PicoFrit fused ilica capillary column (New Objective, Woburn, MA) of 15 cm long was packed with the C18 Jupiter 5 *μ*m 300 Å reverse-phase material (Phenomenex, Torrance, CA). This column was installed on the Easy-nLC II system (Proxeon Biosystems, Odense, Denmark) and coupled to the LTQ Orbitrap Velos (ThermoFisher Scientific, Bremen, Germany) equipped with a Proxeon nanoelectrospray ion source. The buffers used for chromatography were 0.2% formic acid (buffer A) and 100% acetonitrile/0.2% formic acid (buffer B). During the first 12 minutes, 5 *μ*L of sample were loaded on column at a flow rate of 600 nL/ minutes and, subsequently, the gradient went from 2–80% buffer B in 60 minutes at a flow rate of 250 nL/ minutes and then came back at 600 nL/ minutes and 2% buffer B for 10 minutes. LC-MS/MS data acquisition was accomplished in positive ion mode using a eleven scan event cycle comprised of a full scan MS for scan event 1 acquired in the Orbitrap. The mass resolution for MS was set to 60,000 (at m/z 400) and used to trigger the ten additional MS/MS events acquired in parallel in the linear ion trap for the top ten most intense ions. Mass over charge ratio range was from 360 to 2000 for MS scanning with a target value of 1,000,000 charges and a maximum inject time of 100 ms. To improve the mass accuracy, the lock mass option was enabled and decamethylcyclopentasiloxane (m/z 371.101233) was used for real time internal mass calibration. Mass over charge ratio range for MS/MS scanning was from ~1/3 of parent m/z ratio to 2000 with a target value of 10,000 charges and a maximum inject time of 100 ms. The data dependent scan events used a maximum ion fill time of 100 ms and 1 microscan. Target ions already selected for MS/MS were dynamically excluded for 25 s after two MS/MS acquisitions. Nanospray and S-lens voltages were set to 1.3–1.7 kV and 50 V, respectively. Capillary temperature was set to 225°C. MS/MS conditions were: normalized collision energy, 35 V; activation q, 0.25; activation time, 10 ms. The minimum signal required to trigger an MS/MS event was set to 2000 and the isolation width for MS/MS scanning was set to 2. The peak list files were generated with extract_msn.exe (version January 10, 2011) using the following parameters: minimum mass set to 600 Da, maximum mass set to 6000 Da, no grouping of MS/MS spectra, precursor charge set to auto, and minimum number of fragment ions set to 10. Protein database searching was performed with Mascot 2.3 (Matrix Science) against the human NCBInr protein database (version July 18, 2012). The mass tolerances for precursor and fragment ions were set to 10 ppm and 0.6 Da, respectively. Trypsin was used as the enzyme allowing for up to 2 missed cleavages. Mono, di- and tri-methylation of arginine were allowed as variable modifications while carbamidomethylation was set as a fixed modification.

The mascot score cutoff used for the MS analysis were based on the following values:

| Protein score | Percentage of false positive |
| --- | --- |
| >50 | <1 |
| 40-50 | 10 |
| 30-40 | 40 |
| 20-30 | 60 |
| <20 | >90 |

The relative abundance of indicated peptides in Figure 4 were estimated from the peak area of reconstructed ion chromatograms obtained with Qual Browser (Xcalibur, Thermo Fisher Scientific) using a m/z tolerance of 10 ppm.

### Proteinase K protection assay

Mitochondria from control and LONP1 knocked-down cell lines were resuspended in HIM (200 mM mannitol, 70 mM sucrose, 10 mM HEPES, 1 mM EGTA, adjusted to pH 7.5 with KOH) buffer at 1 mg/ml concentration containing the indicated proteinase K concentration (Figure S3). Samples were incubated for 20 minutes on ice. Addition of 2 mM PMSF (phenylmethylsulfanoxide) for 20 minutes stopped the reaction. Samples were spun down for 20 minutes at 20 000 g. The pellet fraction was resuspended in 50 μl of 1X laemmli buffer, sonicated and loaded onto a 12% acrylamide:bisacrylamide (29:1) gel. Subsequent immunoblotting was done with indicated antibodies.

### 2D-SDS PAGE analysis

Mitoplasts were prepared from fibroblasts, as described previously (39) by treating the cells with 0.8 mg digitonin/mg of protein. Mitoplasts were solubilized with 1% n-Dodecyl-β-D-Maltoside, and solubilized proteins were used for electrophoresis. BN-PAGE (40) was used to separate the samples in the first dimension on 6–15% polyacrylamide gradients. For analysis of the second dimension, strips of the first-dimension gel were incubated for 30 minutes in 1% SDS and 1% -mercaptoethanol. The strip was then run on 10% tricine/SDS-PAGE. Individual structural subunits of complexes I and IV were detected by immunoblot analysis as indicated.

### Electron microscopy

Control and LONP1 knocked-down cell lines were incubated overnight at 4°C in fixation solution containing 2.5% glutaraldehyde, 0.1 M sodium cacodylate and 4% sucrose. Next, samples were incubated in post-fixative solution containing 1% osmium tetraoxide and 1.5% potassium ferrocyanide, followed by 15 minutes incubations in increasing concentrations of ethanol (30%-100%) to dehydrate; next samples were infiltrated for 30 minutes with epon/ethanol 1:1, then 1:3 and finally with 100% epon. Samples were embedded in fresh epon for 48 hours at 60 °C, cut and analyzed by transmitted electron microscopy.

### Quantitative RT-PCR

RNA was purified from control and LONP1 siRNA knock-down fibroblasts cell lines (MCH64) using the RNeasy Plus Kit (QIAGEN). cDNA was obtained from 2 μg of purified RNA using the High-capacity cDNA Reverse Transcription kit (4368814) from Applied Biosystems. qPCR was performed using the KAPA SYBR FAST qPCR master mix and run on a Quant Studio 6 Flex system. qPCR analysis was done by Absolute Quantification/2^nd^ derivative of three independent biological replicates, each performed in triplicate. The statistical analysis of mRNA transcript abundance was done after normalization with GAPDH. The statistics software Graphpad Prism 6 was used to performed a two-way ANOVA with Bonferrioni multiple comparisons. Error bars represent standard error of the mean.

ATF4-F: CAG CAA GGA GGA TGC CTT CT

ATF4-R: CCA ACA GGG CAT CCA AGTC

CHAC1-F: GTG GTG ACG CTC CTT GAA GA

CHAC1-R: TTC AGG GCC TTG CTT ACC TG

ASNS-F: GATGAACTTACGCAGGGTTACA

ASNS-R: CACTCTCCTCCTCGGCTTT

PCK2-F: AAACCCTGGAAACCTGGTG

PCK2-R: CAATGGGGACACCCTCTG

ATF5-F: GAG CCC CTG GCA GGT GAT

ATF5-R: CAG AGG GAG GAG AGC TGT GAA

CLPP-F: AAG CAC ACC AAA CAG AGC CT

CLPP-R: AAG ATG CCA AAC TCC TGGG

HSPA9-F: TGG TGA GCG ACT TGT TGG AAT

HSPA9-R: ATT GGA GGC ACG GAC AAT TTT

HSPD1-F: ACTCGGAGGCGGAAGAAA

HSPD1-R: TGTGGGTAACCGAAGCATTT

GAPDH-F: AGGGTCATCATCTCTGCCCCCTC

GAPDH-R: TGTGGTCATGAGTCCTTCCACGAT

## Authors contribution

OZR and EAS conceived the project and designed experiments. OZR executed all experiments and analyzed the data.

## Acknowledgements

This work was supported by a CIHR grant to EAS. O.Z.R. was supported by fellowships from CONACyT (Consejo Nacional de Ciencia y Tecnologia) (209378) and PBEEE (Fonds quebecois de la recherche sur la nature et les technologies) (140802). We thank Woranontee Weraarpachai for reagents and support, Neil Webb for the generation of the LCD, catalytic dead LONP1 construct and Denis Faubert from IRCM for the generation of the mass spectrometry data and MS quantitative analysis.

